# The minimal behavioral time window for reward conditioning in the nucleus accumbens of mice

**DOI:** 10.1101/641365

**Authors:** Kenji Yamaguchi, Yoshitomo Maeda, Takeshi Sawada, Yusuke Iino, Mio Tajiri, Ryosuke Nakazato, Haruo Kasai, Sho Yagishita

**Affiliations:** Laboratory of Structural Physiology, Center for Disease Biology and Integrative Medicine, Faculty of Medicine; International Research Center for Neurointelligence (WPI-IRCN), UTIAS, The University of Tokyo, Bunkyo-ku, Tokyo, Japan

**Author notes:** These authors contributed equally to this work (Kenji Yamaguchi, Yoshitomo Maeda). Corresponding authors: Haruo Kasai, M.D., Ph.D. and Sho Yagishita, M.D., Ph.D., Laboratory of Structural Physiology, Center for Disease Biology and Integrative Medicine, Faculty of Medicine, The University of Tokyo, Faculty of Medicine Bldg.1 #NC207, 7-3-1 Hongo, Bunkyo-ku, Tokyo 113-0033, JAPAN, and.

## Abstract

The temporal precision of reward-reinforcement learning is determined by the minimal time window of the reward action—theoretically known as the eligibility trace. In animal studies, however, such a minimal time window and its origin have not been well understood. Here, we used head-restrained mice to accurately control the timing of sucrose water as an unconditioned stimulus (US); we found that the reinforcement effect of the US occurred only within 1 s after a short tone of a conditioned stimulus (CS). The conditioning required the dopamine D1 receptor and CaMKII signaling in the nucleus accumbens (NAc). The time window was not reduced by replacing CS with optogenetic stimulation of the synaptic inputs to the NAc, which is in agreement with previous reports on the effective dopamine timing of NAc synapses. Thus, our data suggest that the minimal reward time window is 1 s, and is formed in the NAc.

## Introduction

Animal behaviors are effectively reinforced when a reward shortly follows a preceding sensorimotor event for both operant (Thorndike, 1911) and classical (Pavlov, 1927) conditioning, indicating that reward timing is critical for learning. In reinforcement learning theory (Sutton and Barto, 1998; Gerstner et al., 2018), the minimal time window during which a reward can effectively reinforce preceding events is called the eligibility trace which sets a tradeoff between the sensitivity and precision of reward detection: A longer eligibility trace enables the sensitive detection of a delayed reward, but reduces the precision in the detection of a temporal contiguity. How the brain mechanism resolves this tradeoff, however, has not been determined since previous behavioral studies largely focused on long delay conditioning (Boice and Denny, 1965; Holland, 1980; Rescorla, 1988) and the actual minimal time window in behavior and its origin have never been investigated.

Dopamine mediates reward-related signals in mammalian brains. Unexpected rewards cause a phasic burst firing of dopamine neurons in the midbrain, whose activity is reduced after the acquisition of reward conditioning. It has been therefore suggested that dopamine encodes a reward prediction error (RPE) signal (Schultz et al., 1997). The nucleus accumbens (NAc) is a major projection site of dopamine neurons (Menegas et al., 2017), and receives glutamatergic inputs from several brain regions including the amygdala and the hippocampus, showing transient activation to reward predictive cues (Reed et al., 2018). The convergent dopaminergic and glutamatergic signals regulate reward conditioning through *N*-methyl-D-aspartate type glutamate receptors (NMDARs) and dopamine D1 receptors (D1Rs) in the NAc (Kelley et al., 1997; Smith-Roe and Kelley, 2000). In slice preparations, NMDAR, Ca^2+^/calmodulin–dependent protein kinase II (CaMKII) and D1R regulate the enlargement of the dendritic spine, a structural basis for long-term potentiation of the D1R-expressing spiny projection neurons (D1-SPNs) in the NAc (Yagishita et al., 2014). Of note, the phasic burst of dopamine potentiated the spine enlargement when dopamine followed glutamatergic inputs within 0.3 – 1 s in the D1-SPNs (Yagishita et al., 2014) and similar narrow time windows of dopamine for plasticity have been shown in other studies (Wieland et al., 2015; Fisher et al., 2017). The behavioral time window, however, can also be affected by the times for sensory and reward information processing, and by other rules for plasticity (Hebb, 1949; Cassenaer and Laurent, 2012; Fremaux and Gerstner, 2015; Brea et al., 2016).

We therefore systematically characterized the minimal time window of reward conditioning and its origin by developing a conditioning system in head-restrained mice where a tone (conditioned stimulus, CS) was followed by the water reward (unconditioned stimulus, US) which induced licking as unconditioned reflex (UCR). This tone-water-licking task enabled to precisely present US without any delay in consumption and to rapidly establish classical conditioning within an hour, in contrast to tasks where licking is reinforced by water (antecedent-licking-water operant conditioning), which require several days for their acquisition (Sippy et al., 2015). We first investigated the optimal tone duration for conditioning and then characterized the minimal time window. We applied a local infusion of a D1R antagonist and the viral delivery of a CaMKII inhibitory peptide into the NAc to determine the brain regions responsible and the molecular underpinnings. Finally, we used optogenetic stimulation of NAc to eliminate the possible delay of the sensory stimulus to the NAc, and to test the involvement of NAc in CR.

## Results

### Time window for classical conditioning in head-restrained mice

We used a head-restrained system to deliver a US of water at an arbitrary timing for classical conditioning. The position of the licking port was set close to the mouth of the mice (Fig. 1a), so that a drop of water would immediately touch the mouse to signify delivery of the US. Thus, licking responses (UCR) were induced just after the presentation of the US (Fig. 1b). Before conditioning, we measured baseline responses to a short, pure tone (8 kHz, 0.5 s) (Fig. 1c), which was subsequently used as the CS, and confirmed that the tone itself did not evoke a licking response (Fig. 1d). For the tone-water-licking conditioning, we first presented a CS followed by a US at the CS offset for 180 trials (Fig. 1e, f). To monitor the formation of the association during conditioning, 20 CS only trials were pseudo-randomly inserted among the 180 trials with CS-US presentation so that 2 CS only trials were included in every 20 trials. The learning curve of the conditioning was obtained by plotting the lick scores calculated using the averaged licking frequency for the period from the onset of CS to 1 s after the offset, which was subtracted from the lick frequency 2 s before CS (Fig. 1g). The results showed that mice started to predict US arrival at the presentation of the CS after 40 trials of pairing and learning was saturated after 120 trials (Fig. 1g, one-way repeated-measurement ANOVA, F(10, 76) = 8.1, *P* = 4.1 × 10^−8^, post-hoc test by Bonferroni: Baseline vs. 1–20, *P* = 1.0; vs. 21–40, *P* = 1.0; vs. 41–60, *P* = 0.024; vs. 61–80, *P* = 0.0001; vs. 81–100, *P* = 0.0002; vs. 101–120, *P* = 0.0985; vs. 121–140, *P* = 0.0010; vs. 141–160, *P* = 0.0009; vs. 161–180, *P* = 0.0007; vs. 181–200, *P* = 0.003).

**Figure 1.**
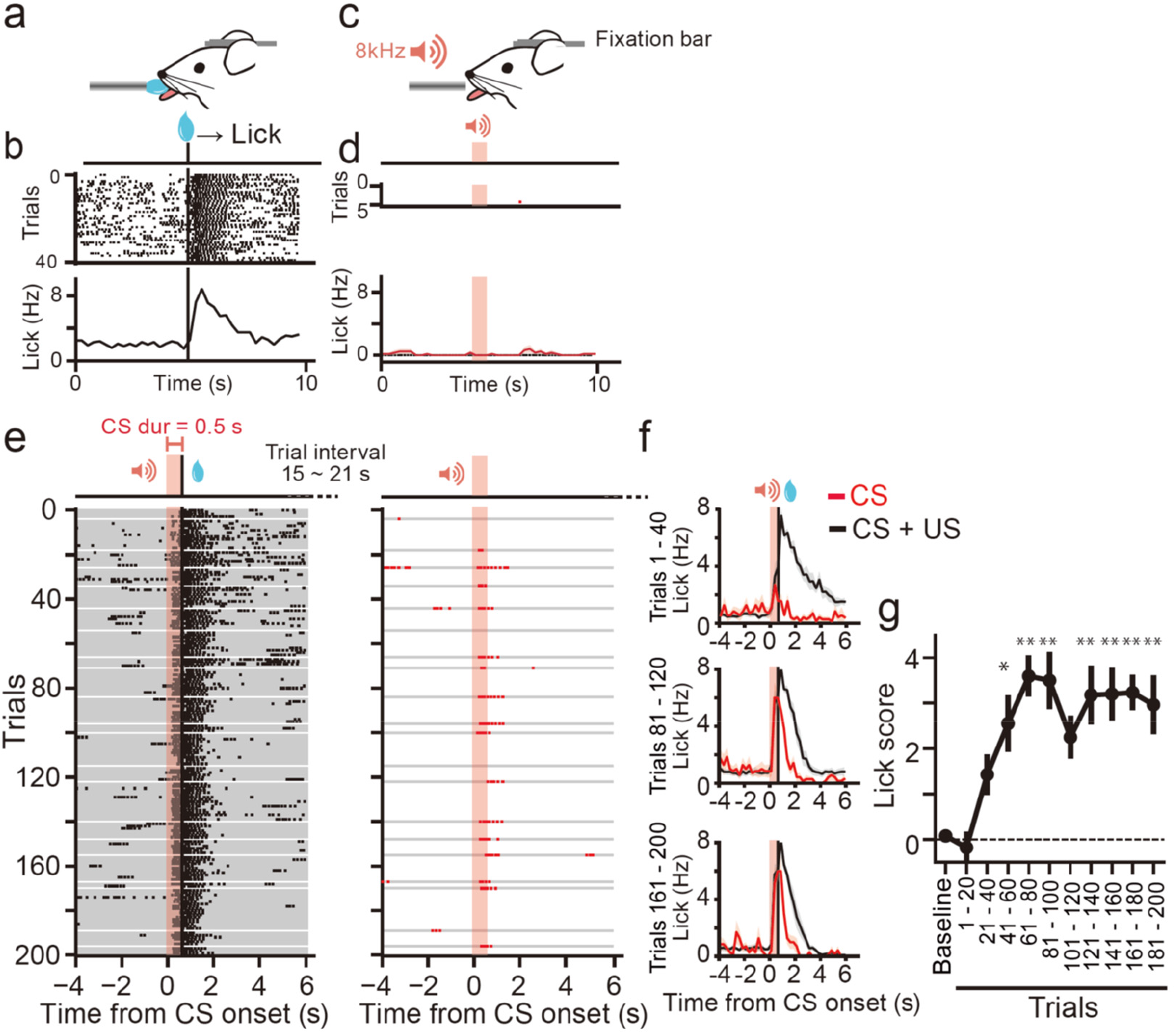
Pavlovian conditioning in head-restrained mice. a) A schematic diagram of the behavioral setup. Mice were head-restrained using a chronically implanted bar. A drop of glucose water was presented as an US to directly touch the mouth of the mouse from a tube that monitored the licking responses. b) Raster plots (upper) and an averaged peri-stimulus time histogram (PSTH, bottom) for the licking responses to US presentation from a representative mouse. The US was presented 40 times at 10-s intervals. Black vertical bars indicate the onset of US presentation. c) A short tone (8 kHz, 70 dB, 0.5 s) presentation without US. d) Raster plots (upper) and an averaged PSTH (bottom, *n* = 7 mice) for licking responses to CS before conditioning. The baseline measurement consisted of five consecutive trials with CS-only presentation. Red shades indicate the period of CS presentation. e) Raster plots for licking during conditioning with a CS duration of 0.5 s in a representative mouse. The session consisted of 180 trials with a paired presentation of CS and US (left) and pseudo-randomly inserted 20 trials with CS-only presentation (right). “CS dur” indicates the duration of CS presentation. US was presented at the offset of CS. Intervals between CS presentations were pseudo-randomly varied between 15 and 21 s. Each gray line indicates a single trial. f) Averaged PSTHs for the licking responses in the first 20% of the trials (1–40, upper), the third 20% of the trials (81 – 120, middle), or the last 20% of the trials (161 – 200, bottom) during CS + US trials (black trace) or CS-only trials (red trace). In the CS-only trials, the average of four trials were included in the indicated periods are shown. The shadows indicate SEM. *n* = 7 mice. g) Lick scores (Materials and Methods) plotted against time. * *P* < 0.05, ** *P* < 0.01 (*n* = 7 mice, one-way ANOVA with repeated measurement). Error bars indicate SEM.

We then examined the dependence of the conditioning on the duration (0.2 s, 0.5 s, 1 s, 2 s, 3 s, and 4 s) of the CSs which were applied at the offset of the CSs (Fig. 1-figure supplement 1). CS durations of 0.5 s and 1 s were effectively reinforced but durations of 0.2 s or longer than 2 s were less effective. A learning curve was obtained by plotting the lick scores during the conditioning (Fig. 1-figure supplement 1g). These results showed that the shortest duration of CS for effective conditioning was 0.5 s (Fig. 1-figure supplement 1h, two-way repeated measurement ANOVA, CS duration × conditioning periods with the latter as a within-subject factor; effect of CS duration: F (1, 63) = 36.4, *P* = 2.0 × 10^−6^; effect of conditioning periods: F (5, 63) = 5.3, *P* = 0.0017; interaction: F(5, 63) = 4.1, *P* = 0.0070; main effect: CS duration = 0.2 s, *P* = 0.13; CS duration = 0.5 s, *P* = 1.6 × 10^−6^; CS duration = 1 s, *P* = 0.00028; CS duration = 2 s, *P* =0.0054; CS duration = 3 s, *P* = 0.52; CS duration = 4 s, *P* = 0.74).

Since the most effective CS duration was determined to be 0.5 s, we determined the minimal time window of the reward action by presenting US with various delays (Fig. 2a-f). When the US preceded the CS, the CS did not induce licking responses after conditioning (Fig. 2a-b). When the CS preceded the US by no more than 1 s, the mice rapidly predicted the US (Fig. 2c-e). However, a CS-US interval of 2-s did not allow the formation of the association (Fig. 2f). For this analysis, the lick scores were calculated from the averaged licking frequency for 2 s after CS presentation subtracted from that 2 s before CS presentation to plot a learning curve (Fig. 2g) and time window (Fig. 2h). We found that the minimal time window of conditioning was 1 s (Fig. 2h) (two-way repeated measurement ANOVA, US delay × conditioning periods with the latter as a within-subject factor; effect of US delay: F (1, 79) = 20.7, *P* = 6.6 × 10^−5^; effect of conditioning periods: F (5, 79) = 8.0, *P* = 4.4 × 10^−5^; interaction: F(5, 79) = 6.3, *P* = 0.00032; main effect: Δt = −1, *P* = 0.99; Δt = −0.5, *P* = 0.72; Δt = 0, *P* = 0.42; Δt = +0.5, *P* = 8.6 × 10^−6^; Δt = +1, *P* = 4.0 × 10^−7^; Δt = +2, *P* = 0.77).

**Figure 2.**
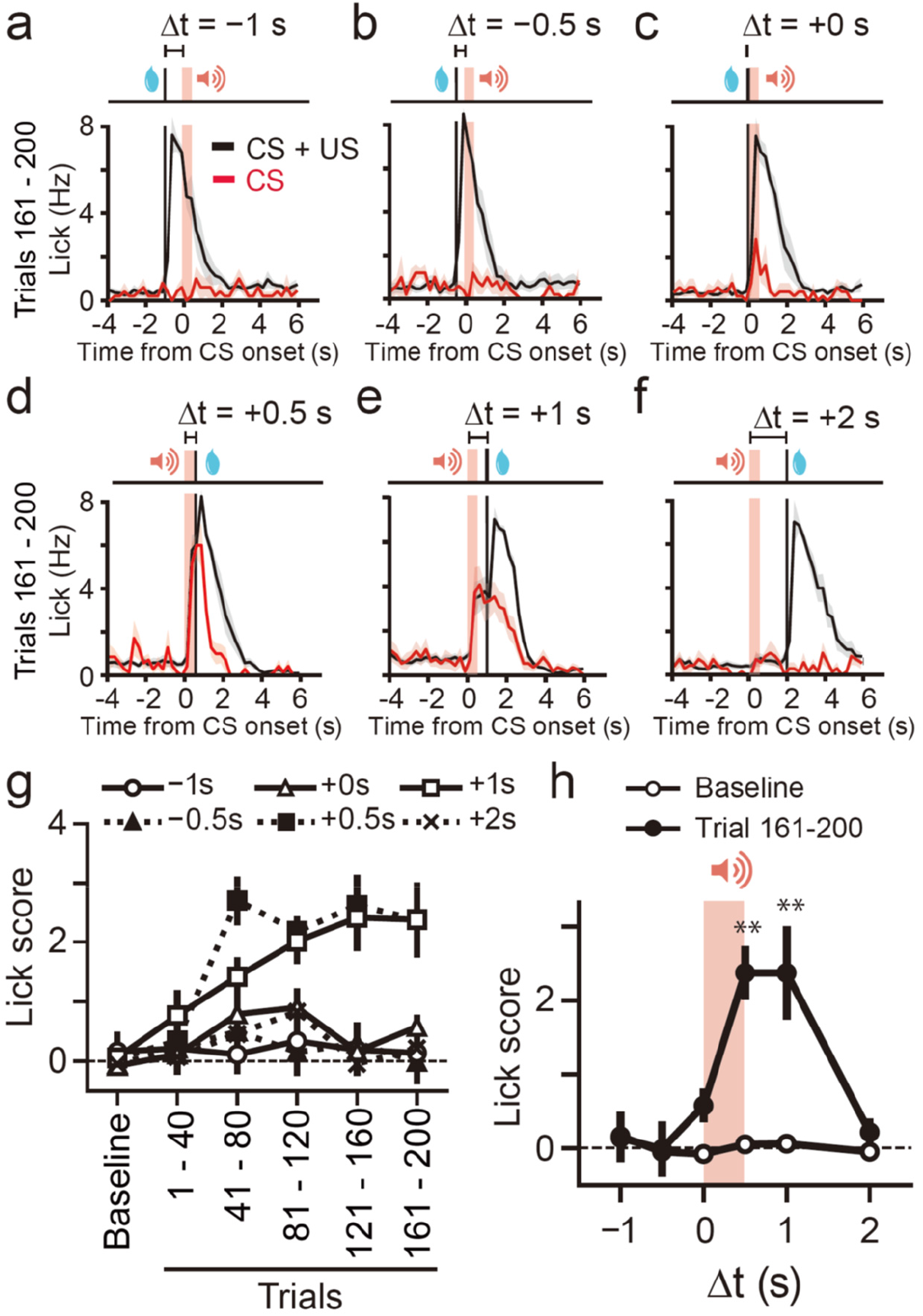
Conditioning with various delays of the US. a-f) Averaged PSTHs of the licking responses with delays of −1 s (a, *n* = 5 mice), −0.5 s (b, *n* = 5 mice), +0 s (c, *n* = 5 mice), +0.5 s (d, *n* = 7 mice), +1 s (e, *n* = 11 mice), or +2 s (f, *n* = 7 mice) during trials 161–200. The plot in (d) is the same as that in the bottom of Fig. 1f. Red shades indicate the period of CS presentation. g) Peak lick scores plotted against time. Each symbol represents the delay in the US relative to the CS.Time windows for US presentation leading to conditioning. The averaged lick scores in the baseline session (white circle) and CS-only trials included during conditioning trials 161–200 (black circle) were plotted against delays between the CS and US. Two-way ANOVA with the test period as the repeated measurement factor. ** *P* < 0.01.

### Role of the NAc in Pavlovian conditioning with the shortest CS duration

We tested whether the molecular signaling required for plasticity in the NAc is responsible for the shortest form of conditioning. First, we injected a dopamine D1R antagonist (SCH23390) in the bilateral NAc during conditioning (Fig. 3a). A D1R antagonist blocked the conditioning when the CRs were measured at the end of conditioning (Fig. 3b-e) (t-test, t(8) = 2.38, *P* = 0.044). The D1R antagonist also partially inhibited US responses, suggesting that D1R inhibition affected motor components. However, CRs on the next day where no drug was present were also inhibited in mice with the D1R antagonist (Fig. 3f).

**Figure 3.**
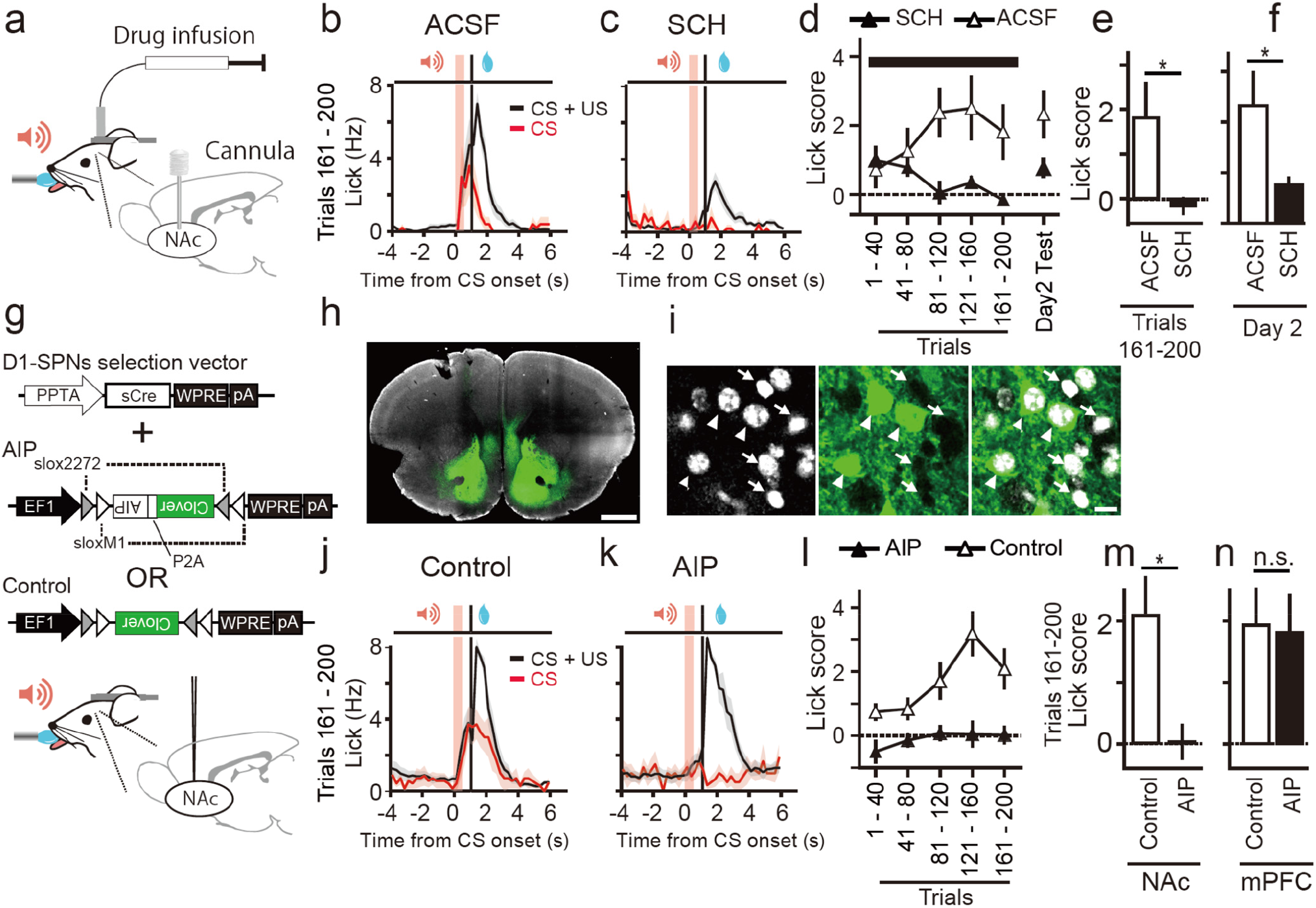
The effect of a D1 receptor blocker and CaMKII inhibitory peptide (AIP) in the NAc on conditioning. a) A schematic diagram for drug infusion into the NAc. b, c) Averaged PSTHs of the licking responses in trials 161-200 with injection of a D1 blocker, ACSF (b, *n* = 5 mice) or SCH23390, (c, *n* = 5 mice) during conditioning. d) The lick scores plotted against time for the two conditions indicated. Drug infusion was started 30 min before the behavioral experiments and was continued during conditioning as indicated by the black horizontal bar. e, f) The effect of a dopamine D1 receptor blocker, SCH23390, on conditioning. Averaged lick scores of CS-only trials during trials 161–200 (e) or day 2 (f) were plotted. * *P* < 0.05, t-test, *n* = 5 mice for SCH23390, *n* = 5 mice for ACSF. g) Viral constructs and schematics of injection. h, i) Confocal images of clover fluorescence (green) and DAPI (white) from a coronal slice including the NAc. Slices were counter stained with DAPI, and 35.8 % (72/201) of cells were positive for AIP. Arrow heads indicate AIP-positive cells and arrows indicate negative cells. Scale bars indicate 1 mm (h) and 20 μm (i). j, k) Averaged PSTHs of the licking responses in CS + US (black) and CS (red) conditionings from mice injected with control (j, *n* = 7 mice) or AIP (k, *n* = 6 mice). l) The peak lick scores plotted against time for the conditions indicated. m, n) Average lick scores from eight trials during trials 161–200 from mice with injections in the NAc or injections in the mPFC (Figure3-figure supplement 2) were plotted for each condition. * *P* < 0.05, t-test.

Then, we examined whether CaMKII is also required for this conditioning. For this purpose, we developed an AAV vector delivering autocamtide 2-related inhibitory peptide (AIP), a peptide that blocks CaMKII, in D1-SPNs using a PPTA promoter (Fig. 3g). AAV was injected bilaterally into the NAc and the extent of the expression was monitored by a green fluorescent protein that was co-expressed with AIP using a P2A cleavage site (Fig. 3h, i). We first confirmed whether virally-delivered AIP blocked the structural plasticity of SPNs in acute slices from the NAc. To test this, we used Mg-free artificial cerebrospinal fluid (ACSF) which causes a strong influx of calcium that induced spine enlargement by two-photon uncaging of caged glutamate, even in the absence of dopamine in the control vector expressing SPNs (Fig. 3-figure supplement-1a-c). With this strong protocol, AIP-expressing neurons showed inhibition of the spine enlargement at the sustained phase compared to the control group (Fig. 3-figure supplement-1b-e) (t-test, t(10) = 3.4, *P* = 0.0067), suggesting that AIP effectively inhibited structural plasticity in the NAc, similarly to the hippocampus (Murakoshi et al., 2017). Although AIP may have affected the cellular signaling, the early phase of the structural plasticity was not affected as in the case of acute pharmacological inhibition (Matsuzaki et al., 2004).

Having confirmed the inhibitory effects of AIP, we tested the behavioral effects of AIP expression in the NAc and found that that the expression of AIP in the NAc abolished learning (Fig. 3j-m) (t-test, t(11) = 2.7, *P* = 0.019). In contrast, expression of AIP in the prefrontal cortex under a CaMKII promoter did not affect conditioning (Fig. 3n, Fig. 3-figure supplement-2)(t-test, t(10) = 0.14, *P* = 0.89). These results indicated that reward conditioning with a short CS duration preferentially relied on the NAc molecular signaling related to plasticity.

### Optogenetic stimulation of the synaptic input to the NAc for conditioning

Although natural tone of the CS showed that 1 s was the shortest time window, it is still possible that the observed window was prolonged by the neuronal processes that carry the sensory signal to the NAc. To exclude this possibility, we applied optogenetics to directly stimulate glutamatergic inputs to the NAc. We stimulated axons from the basolateral amygdala (BLA) because previous studies showed that the BLA to NAc circuit is involved in cue-reward association (Gallagher et al., 1990; Stuber et al., 2011). The ChR2-expressing AAV vector was injected into the left amygdala and an optical fiber was placed in the ipsilateral NAc (Fig. 4a, b). First, we replicated the previous findings indicating that the stimulation of the axonal projection from the BLA to the NAc reinforced operant behavior (Fig. 4-figure supplement-1a-b)(Stuber et al., 2011) by stimulating axonal fibers (457 nm, 5 ms, 20 Hz, 10 times) at high (> 5 mW) laser power (Fig. 4-figure supplement-1c, d) (one-way repeated measure ANOVA, F(3, 45) = 86.0, *P* = 1.2 × 10^−18^, Bonferroni post-hoc analysis, laser on at low power vs. laser off, *P* = 1.0, laser on at high power vs. laser off, *P* = 1.2 × 10^−16^, laser on at low power vs. laser on at high power, *P* = 2.4 × 10^−15^). In contrast, subthreshold low laser powers (< 3 mW) did not reinforce this behavior (laser on at low power vs. laser off at low power, *P* = 1.0) (Fig. 4-figure supplement-1c, d).

**Figure 4.**
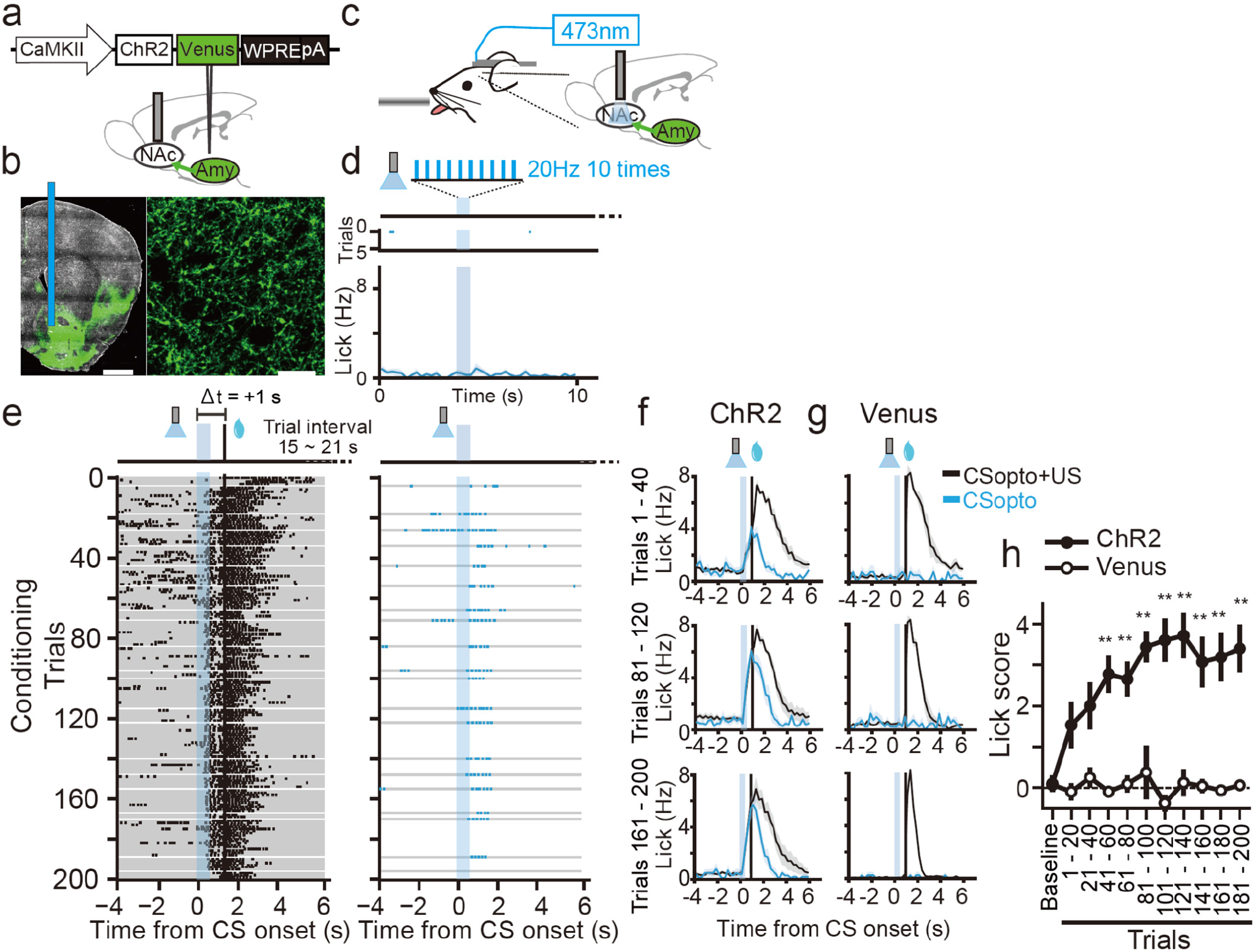
Pavlovian conditioning with CSopto. a) A viral construct and schematic of the AAV injection and optical fiber implantation. ChR2-expressing AAV was injected into the left basolateral amygdala (BLA). An optical cannula (200 μm core) was placed into the left NAc. b) Macroscopic (left) and microscopic (right) confocal images of the green fluorescence of Venus fused with ChR2 in the NAc. A blue vertical bar indicates the tract of the inserted optical cannula. Scale bars indicate 1 mm (left) and 20 μm (right). c) Schematic of the behavioral setup. The optical cannula was connected to a laser (473nm) by a patch cable. d) Representative licking responses before conditioning in response to ChR2 stimulation (CSopto, 20 Hz, 10 times, 5 ms pulse width). Raster plots indicating licking responses from a representative mouse and PSTHs indicating averaged responses over 10 mice. Blue shades indicate the period of CSopto presentation. e) Representative licking responses during CSopto conditioning with Δt = +1 s. The conditioning paradigm was the same as Fig. 1 except that the tone was replaced with CSopto and the delay was 1s. Raster plots indicating licking responses from CS+US trials (left) or CS-only trials (right). Each gray bar indicates a single trial. Black vertical bars indicate the onset of US presentation. f, g) Averaged PSTHs for the licking responses in the first 20% of the trials (1–40, upper), the third 20% of the trials (81–120, middle), or the last 20% of the trials (161–200, bottom) for each of the CS + US trials (black trace) or the CS-only trials (blue trace) from mice injected with ChR2 (f, *n* = 10) or Venus (g, n = 4). Shadows indicate SEM. h) Lick scores (Materials and Methods) plotted against time course for the ChR2 mice (*n* = 7) and Venus mice (n = 4). ** *P* < 0.01. Error bars indicate SEM.

We then tested whether this weak stimulation of synaptic inputs (optogenetic conditioned stimulus, CSopto) could be associated with the US. In head-fixed mice, blue light stimulation (20 Hz, 0.5 s, 5 ms pulse) of CSopto alone in the NAc did not cause the licking response (Fig. 4c, d). When CSopto was paired with a US of water (Fig. 4e-f), the mice started to show anticipatory licking to CSopto within 30 trials (Fig. 4e, f, h, one-way repeated-measurement ANOVA, F(10, 109) = 6.8, *P* = 7.0 × 10^−8^, post-hoc test by Bonferroni: Baseline vs. 1–20, *P* = 0.92; vs. 21–40, *P* = 0.095; vs. 41–60, *P* = 0.009; vs. 61–80, *P* = 0.0021; vs. 81–100, *P* = 9.2 × 10^−6^; vs. 101–120, *P* = 2.7 × 10^−6^; vs. 121–140, *P* = 1.2 × 10^−6^; vs. 141–160, *P* = 0.0001; vs. 161–180, *P* = 0.0001; vs. 181–200, *P* = 1.1 × 10^−5^). In contrast, mice injected with a Venus vector without ChR2 did not form an association (Fig. 4g, h) (F(10, 43)=0.59, *P* = 0.81), indicating that mice did not respond to optical stimulation itself as a CS but the conditioning relied on optically induced synaptic activation. Moreover, CSopto conditioning was dependent on the D1R, which was tested using a within-subject design to functionally confirm virus injection and fiber placement for ChR2 excitation (Fig. 4-figure supplement-2, paired t-test, t(8) = 3.2, *P* = 0.012).

Finally, we examined the time window of the association formed by the CSopto (20 Hz, 0.5 s) (Fig. 5). The time window of conditioning by the CSopto was within 1 s after the onset of CSopto (Fig. 5h) (two-way repeated measurement ANOVA, US delay × conditioning periods with the latter as a within-subject factor, effect of US delay, F (1, 71) = 16.7, *P* = 0.00029; effect of conditioning periods, F(5, 71) = 9.5, *P* = 1.5 × 10^−5^; interaction, F(5, 71) = 7.0, *P* = 0.00018; main effect: Δt = −1, *P* = 0.24; Δt = −0.5, *P* = 0.86; Δt = 0, *P* = 0.060; Δt = +0.5, *P* =0.0031; Δt = +1, *P* = 6.8 × 10^−8^; Δt = +2, *P* = 0.37), similar to the natural tone (Fig. 2h). For those negative conditions (−1s, −0.5 s, 0 s, and 2 s), we confirmed successful conditioning with 1 s delay on the next day (Fig. 5-figure supplement-1), indicating that the negative results were not due to inappropriate virus injection nor optical fiber placement.

**Figure 5.**
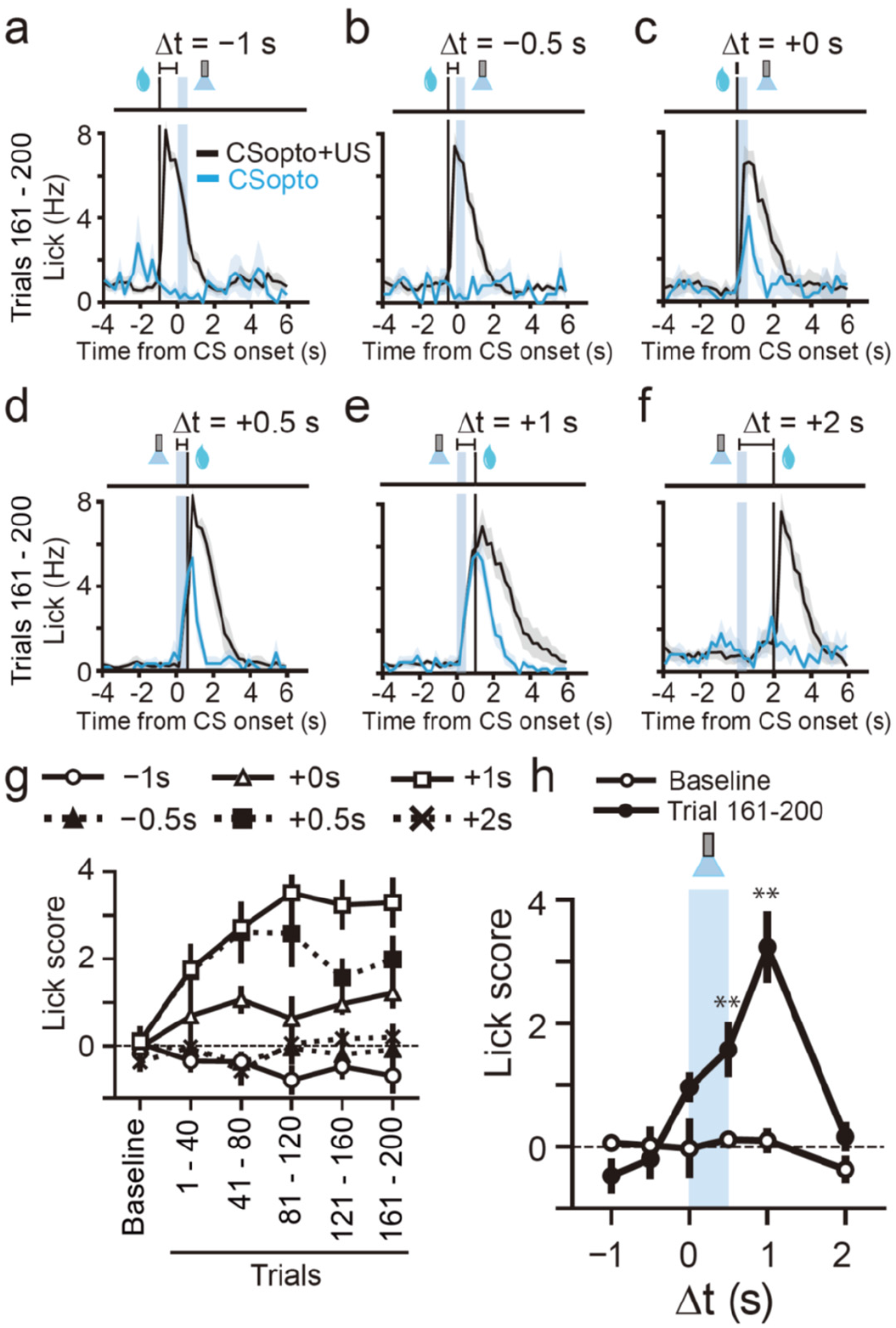
Various delays in US for CSopto conditioning. a-f) Averaged PSTHs of the licking responses in conditioning with delays of Δt = −1 s (a, *n* = 5 mice), Δt = −0.5 s (b, *n* = 5 mice), Δt = +0 s (c, *n* = 5 mice), Δt = +0.5 s (d, *n* = 6 mice), Δt = +1 s (e, *n* = 10 mice), and Δt = +2 s (f, *n* = 5 mice) during 161-200 trials. (e) is the same plot as the bottom trace of Fig. 4f. g) Lick scores plotted against time course for each condition. h) Time window for conditioning. Lick scores in test trials or eight trials during trials 161–200 were plotted against delays between the CS and US. Two-way ANOVA with test period as the repeated measurement factor. ** *P* < 0.01.

## Discussion

We have developed a tone-water-lick classical conditioning in head-restrained mice, and found that the time window for the conditioning was 1 s. The optogenetic stimulation of glutamatergic inputs in the NAc was enough to form the time window but did not further shorten the window. The conditioning was dependent on D1R and CaMKII in the NAc, and the time window closely matched with dopamine time window for spine plasticity in the NAc (Yagishita et al., 2014), which is formed by D1R, Ca^2+^ priming of adenylate cyclase, PKA and CaMKII activation. These data suggested that the minimal time window of classical conditioning is 1 s, and the window is formed in the NAc. Unlike the synaptic dopamine timing in brain slices, the behavioral time window was marginally positive at 0 s which may be ascribed to the delay in the onset of dopamine firing after the US presentation at about 0.2 s *in vivo* (Schultz et al., 1997; Eshel et al., 2015). Our estimate of 1 s was shorter than a previous estimate of 4 s in freely moving animals (Boice and Denny, 1965), where reward timing was not precisely controlled.

Importantly, the minimal reward time window was longer than the burst duration of dopamine neurons of ~0.3 s (Schultz et al., 1997; Eshel et al., 2015), indicating that the 1 s time window effectively prolongs the duration of reward detection compared to the time scale of dopamine firing, but compromises the precision in detection of the temporal contiguity. This tradeoff may be so critical to the animal that it is determined by molecular timing mechanisms associated with dopamine and Ca^2+^-sensitive adenylate cyclase. Interestingly, similar time windows in orders of seconds for conditioning were found in Aplysia (Hawkins et al., 1986; Abrams and Kandel, 1988) and in insects (Mariath, 1985; Tully and Quinn, 1985; Ito et al., 2008), both of which are dependent on monoamine and Ca^2+^-sensitive adenylate-cyclase. Thus, the time windows for reinforcement learning are longer than the typical temporal scales for the spike-timing mechanisms (~20 ms), and the supervised learning in the cerebellum (~100 ms) (Suvrathan et al., 2016).

We also found that the conditioning efficacy peaked at 0.5 – 1 s from the onset of the CS, in accordance with the dopamine time window that peaked at 0.6 – 1 s from the onset of the glutamatergic inputs in the NAc synapses (Yagishita et al., 2014). These results suggest that the timing mechanism is optimized to detect reward delays for the temporal contiguity rather than the synchronicity of the onsets. The delayed peak may also account for the optimal duration of the tone at 0.5 s: the CS duration of 0.2 s was shorter than the optimal delay; and the longer CS duration was unable to effectively detect the strong activation of glutamatergic inputs that may have peaked soon after the CS onset and which may have waned thereafter (Reed et al., 2018). Along with this, the tuning curve for the tone duration (Fig. 2-figure supplement 1h) showed a similarity with the reward time window (Fig. 2h).

The CSopto experiments suggested that the change in NAc synapses but not upstream sensory systems represented the CS-US association that resulted in licking. After conditioning, the CSopto induced a transient rhythmic licking movement, which was analogous to the licking induced by the US of water, that can be regulated by brainstem pathways downstream of NAc (Roseberry et al., 2016; Rossi et al., 2016). This result suggests that the US signal passes through the NAc to fire neurons for licking (Day et al., 2006), similar to the US pathway in the amygdala for threat conditioning (Maren and Fanselow, 1996; Gore et al., 2015). Thus, it can be envisaged that there are the spiny projection neurons (SPNs) in the NAc that can trigger licking by US, and which receive glutamatergic synaptic inputs during CS. The CS input was not strong enough to trigger licking, but potentiation of the synapses by conditioning caused these SPNs to trigger licking.

Although the time window was never longer than 1 s in our conditioning with the short CS, it could be longer (> 30 s) in other behavioral tasks (Tolman, 1948; Boice and Denny, 1965; Holland, 1980; Bangasser et al., 2006; Moe et al., 2009; Heys and Dombeck, 2018), suggesting that additional mechanisms such as working memory or second-order conditioning may complement the NAc eligibility trace. Short-term memory-like mechanisms or cognitive maps (Wikenheiser and Schoenbaum, 2016) simply explain the long delay learning when persistent glutamatergic inputs are sent to the NAc so that the inputs can be associated with reward using the NAc eligibility trace, as proposed in theoretical studies (Bakker, 2002; Rombouts et al., 2015). In fact, pyramidal neurons in the hippocampus show persistent activity during long-trace reward conditioning (Heys and Dombeck, 2018). Second-order conditioning, where reward predicting CS becomes a reinforcer for further preceding events, also allows the association of longer delays than the NAc eligibility trace (Rescorla and Holland, 1982; Schultz et al., 1997). Alternatively, synaptic mechanisms with longer eligibility traces outside the NAc (Brzosko et al., 2015; He et al., 2015) may play direct roles in more complex reward learning (Tye et al., 2008; Brzosko et al., 2015; Jocham et al., 2016; Otis et al., 2017). Therefore, while the short NAc eligibility trace may become predominant in occasions when reward timing is short such as in odor-food associations or inhalation or injections of addictive substances, other mechanisms may support the temporal flexibility of high-order learning along with the NAc mechanisms. How each particular mechanism for the time window is used in various reward learning, and how the NAc and additional brain mechanisms interplay should also be investigated.

In conclusion, we identified that the minimal time window for reward action was 1 s, which is in close agreement with the dopamine timing of synaptic plasticity in the NAc. Such biologically defined temporal constraints may help to construct neuronally plausible theoretical models for reinforcement learning. Since the behavioral learning tightly reflected the synaptic properties, our conditioning system may provide a way of linking molecular and synaptic function with behavior in disease models.

## Materials and Methods

### Adeno-associated virus (AAV)preparation

We cloned the following AAV-expression plasmids: pAAV-CaMKII(0.3)-hChR2(H134R)-Venus, pAAV-CaMKII(0.3)-Venus, pAAV-PPTA-sCre, pAAV-sDIO(M1)-Clover-P2A-AIP, pAAV-sDIO(M1)-Clover, pAAV-CaMKII(0.3)-mCherry-P2A-AIP and pAAV-CaMKII(0.3)-mCherry. The PPTA promoter, a D1-SPN specific promoter, was cloned from the mouse as described previously (Hikida et al., 2010; Yagishita et al., 2014). Autocamtide 2-related inhibitory peptide (AIP), a CaMKII inhibitory peptide, and self-cleaving 2A peptide of porcine teschovirus-1 (P2A) were fused with Clover and cloned in an sCre dependent double inverted ORF expression vector designed using sloxP and sloxP (M1). The original plasmid containing hChR2(H134R) was a kind gift from Dr. Deisseroth and sCre was purchased from Kazusa DNA Research Institute (Japan) (Suzuki and Nakayama, 2011). AAV vectors were produced and their titers measured as described previously (Grieger et al., 2006). Briefly, plasmids for the AAV vector, pHelper (Stratagene), and RepCap5 (Applied Viromics) were transfected to HEK293 cells (AAV293, Stratagene). After 3 days of incubation, the cells were collected and purified twice using iodixanol. The titers for AAV were estimated using quantitative polymerase chain reaction.

### Animals and surgery

Wild type or DAT-IRES-Cre (B6.SJL-Slc6a3tm1.1(cre)Bkmn/J, The Jackson Laboratory) male B6J mice aged 2–4 months old were used. These mice were housed on a 12-h light/12-h dark cycle. A custom-made titanium plate was attached to the head using dental cement. For AIP experiments in the NAc, a total of 1.5 μl of the AAV mixture of PPTA-sCre (5 × 10^11^ GC/mL) with either EF1-sDIO(M1)-Clover-P2A-AIP (2 × 10^13^ GC/mL) or EF1-sDIO(M1)-Clover (1 × 10^13^ GC/mL) were bilaterally injected (AP +1.3 mm, ML +/− 1.0 mm, DV + 4.5 mm) through a glass pipette. For AIP experiments in the medial prefrontal cortex (mPFC), 1.5 μl of CaMKII (0.3)-mCherry-P2A-AIP (2 × 10^13^ GC/mL) or CaMKII(0.3)-mCherry (2 × 10^13^ GC/mL) were bilaterally injected (AP +1.8 mm, ML +/− 0.3 mm, DV + 2.5 mm). The infusion rate was controlled using a syringe pump set at 0.05-0.1 μl/min. For the ChR2 experiments, 1 μl of CaMKII(0.3)-ChR2-Venus (2–3 × 10^13^ GC/mL) or CaMKII(0.3)-Venus (2–3 × 10^13^ GC/mL) was injected into the left basolateral amygdala (AP −1.6 mm, ML −3.3 mm, DV +4.7 mm). After injection, an optical fiber cannula (200 μm core, 5.0 mm in length, Thorlabs, CFM12) was inserted into the left NAc (AP +1.4 mm, ML − 0.75 mm, DV + 4.1 mm). For the drug infusion experiments, a 5.0 mm double guide cannula (26-gauge, 1.5 mm apart from each cannula, Plastic One) were implanted bilaterally into the NAc (AP +1.3 mm, ML +/− 0.75 mm, DV + 4.2 mm). The experimental protocol was approved by the Animal Experimental Committee of the Faculty of Medicine, The University of Tokyo.

### Behavioral experiments

Mice were allowed 4 days for recovery after head plate installation in experiments without virus injections and 3 weeks for recovery in experiments with virus injections. Mice were then habituated for 3 days to the experimental setup without head fixation and water restricted such that body weight was maintained at no less than 80% of the baseline weight. On the day of the experiment, the mice were head-fixed and the licking responses to tone presentation (8 kHz, 70 dB) used as CS were monitored for 5 trials (day 1, baseline session). For the US, a drop of 5% glucose water (2 μl) was presented through the tip of a lick port controlled by a syringe pump. The position of the lick port was set such that the drop of water contacted the mouth of the mice to induce licking without any training. The conditioning session consisted of 180 trials with presentation of CS-US pairs and 20 trials with presentation of CS only. For the time window experiment, each mouse was assigned to one of the CS-US delays of −1 s, −0.5 s, 0 s, +0.5 s, 1 s or 2 s with CS duration of 0.5 s. For the CS duration experiment, each mouse was assigned to one of the CS durations of 0.2 s, 1 s, 2 s, 3 s, or 4 s. The data from the mice assigned to CS-US delays of +0.5 s were also used as that of the CS duration of 0.5 s. The intervals between the trials were randomized with a uniform distribution between 15 s and 21 s, with a mean of 18 s. To monitor learning during conditioning, CS only trials were pseudo-randomly inserted so that 2 trials with CS only were included in every of 18 CS-US trials to record conditioned reflexes (CRs) without US. The licking responses were electrically measured (Hayar et al., 2006). The control of the stimulus presentations and the recording of the licking responses were performed with custom software written in LabView (National Instruments).

For experiments with ChR2 stimulation, a fiber cannula was connected to a blue laser (473 nm, Thorlabs). For an instrumental session (Stuber et al., 2011), conditions with laser on and off were alternately repeated twice. In the laser on condition, axonal fibers were stimulated (5 ms pulse, 10 times in 20 Hz) 100 ms after the detection of a licking event while no stimulation was made in the laser off condition. After the stimulation, we inserted a 500-ms refractory period for stimulation, even though the sensor detected licking. The number of licking responses were counted for 190 s. To initiate licking, the lick port delivered a drop of water once 10 s before recording. The session was repeated with increasing laser power from 1, 2, 3, 5, 7.5 to 15 mW (200 μm core fiber) or until the mice lick counts during the laser on period were 20 times greater than those during the laser off period. For classical conditioning with ChR2, 20-Hz laser stimulation (5 ms pulse, 1 or 2 mW) given 10 times (CSopto) was substituted for the CS tone.

For the drug infusion experiment, SCH23390 (400 μM, Abcam) dissolved in ACSF (125 mM NaCl, 2.5 mM KCl, 2 mM CaCl_2_, 1 mM MgCl_2_, 1.25 mM NH_2_PO_4_, 26 mM NaHCO_3_, and 20 mM glucose) or ACSF for controls was infused at the rate of 16.66 nl/min by a syringe pump (Legato111, KD scientific) 30 min before the experiments. The infusion was continued during the conditioning at the rate of 14.9 nl/min. For pharmacological experiments during CSopto conditioning, SCH23390 or saline were intraperitoneally injected 30 min before the conditioning experiments. Doses of 0.25 and 0.5 mg/kg were tested. As the results were similar between the doses, the data were pooled in the analysis.

### Histological analysis

For the AIP experiments, the mice were subjected to the histological analysis to confirm AIP expression in the NAc. After the behavioral experiments, the mice were transcardially perfused with 4% paraformaldehyde and decapitated. Coronal slices of 50-μm thickness were obtained. Clover fluorescent was obtained using stereoscopic microscopy (Leica M165-FC), and images were captured with a CMOS camera (Hamamatsu photonics ORCA R2). AIP expression was considered sufficient if it was expressed bilaterally including more than 3/4 of the anterior part of the anterior commissure, a NAc surrounding structure. Out of the 18 NAc injected mice, five failed to satisfy this criterion (one did not exhibit expression at all, three exhibited unilateral expression only, and one exhibited expression only in the medial half of the NAc) and were therefore excluded from behavioral analyses. For some slices, detailed fluorescence images were obtained using confocal microscopy (Leica, SP5) of the preparations which were counter stained using DAPI.

### Slice experiments

Acute coronal slices of the NAc (280 μm) were obtained from 5- to 6-week-old mice injected with either an AIP (PPTA-sCre + sDIO-AIP-clover) or a control vector (PPTA-sCre + sDIO-clover) at 3 weeks of age. Mice were anesthetized using ketamine and xylazine; perfused transcardially with an ice-cold solution containing 220 mM sucrose, 3 mM KCl, 8 mM MgCl_2_, 1.25 mM NaH_2_PO_4_, 26 mM NaHCO_3_, and 25 mM glucose; and then quickly decapitated to prepare slices with a VT1200 microtome (Leica). The slices were incubated at 34°C for 30 min and then at room temperature (20–25°C) in artificial cerebrospinal fluid (ACSF; 125 mM NaCl, 2.5 mM KCl, 1 mM CaCl_2_, 2 mM MgCl_2_, 1.25 mM NaH_2_PO_4_, 26 mM NaHCO_3_, and 20 mM glucose), bubbled with 95% O_2_ and 5% CO_2_. The slices were transferred to a recording chamber and superfused with the same ACSF containing 2 mM CaCl_2_, 1 mM MgCl_2_, and 200 μM Trolox (Sigma) at 30–32°C.

D1-SPNs were identified by somatic expression of clover, and targeted whole-cell recordings were made with a patch-clamp electrode (open-tip resistance, 5—8 MΩ) filled with a solution containing 120 mM potassium gluconate, 20 mM KCl, 10 mM disodium phosphocreatine, 50 μM Alexa 594 (Life Technologies), 4 mM ATP (magnesium salt), 0.3 mM GTP (sodium salt), 10 mM HEPES, and 5 μM β-actin (human platelets; Cytoskeleton) whose pH was adjusted to 7.25 with KOH, and the osmolarity was adjusted to 275—280 mOsm/l with sucrose. The cells were voltage-clamped at −70 mV. Two-photon imaging of the dendritic spines was performed with an upright microscope (BX61WI, Olympus; objective lens, LUMPlanFI/IR, 60×, numerical aperture 1.0) equipped with an FV1000 laser-scanning microscope system connected to two mode-locked, femtosecond-pulse Ti:sapphire lasers (MaiTai from Spectra Physics, 720 nm for uncaging and 980 nm for imaging).

For plasticity induction, Mg-free ACSF was perfused to stimulate NMDA receptors (Matsuzaki et al., 2004) and CDNI-glutamate (Ellis-Davies et al., 2007) (4 mM) was locally pressure-puffed from a glass pipette positioned close to the selected dendrite. The structural plasticity of each single spine was induced using repetitive two-photon uncaging of CDNI-glutamate (5—6 mW, 0.8 ms activation of 720 nm laser) at 10 Hz 200 times in Mg-free ACSF. For each experiment, three to four spines located 100 μm from the soma and 20—35 μm in depth from the slice surface were stimulated. Three-dimensional reconstructions of the dendritic morphology were generated by the summation of fluorescent values separated by 0.5 μm. The fluorescence of the dendrites across the time series was corrected by the entire fluorescence of an imaging area where dendritic fluorescent signals were predominant. Neighboring spines were defined as spines within 3 μm of the stimulated spines.

### Data analysis

For the analysis of the CS-induced licking responses (CRs), we calculated the lick score in the CS-only trials as [average licking frequency (Hz) during 2 s after CS presentation] – [average licking frequency during 2 s before CS presentation] except for the CS duration analysis where the lick frequency from CS onset to 1 s after CS offset was subtracted from the baseline. Analysis of variance (ANOVA) or t-test were adapted for statistical tests with a threshold of *P* < 0.05. The spine-head volumes were estimated from the total fluorescence intensity and the mean volume changes were calculated for each dendrite and the averaged volume changes for each dendrite were subjected to statistical analysis. Data analyses were performed using Excel (Microsoft) and Excel Statistics (SSRI). Data are presented as mean ± SEM.

## Acknowledgements

We thank A. Kurabayashi, M. Asaumi, A. Nishikawa, M. Ikeda for their technical assistance, and S. Ishii for helpful discussion and support. This work was supported by CREST (JPMJCR1652 to H.K.) from JST, SRPBS (JP19dm0107120 to H.K.), BRAIN/MINDS (19dm0207069h0001 to S.Y.) from AMED, Grants-in-Aid (No. 26221001 to H.K.;15K18333, 19K16249, 16H06395, 16H06396 and 16K21720 to S.Y.) from JSPS, the World Premier International Research Center Initiative (WPI) from MEXT, and Takeda Science Foundation (to S.Y.). M.T. and T.S. are the Research Fellows for Young Scientists of JSPS.

---

**Figure 1-Figure supplement-1.**
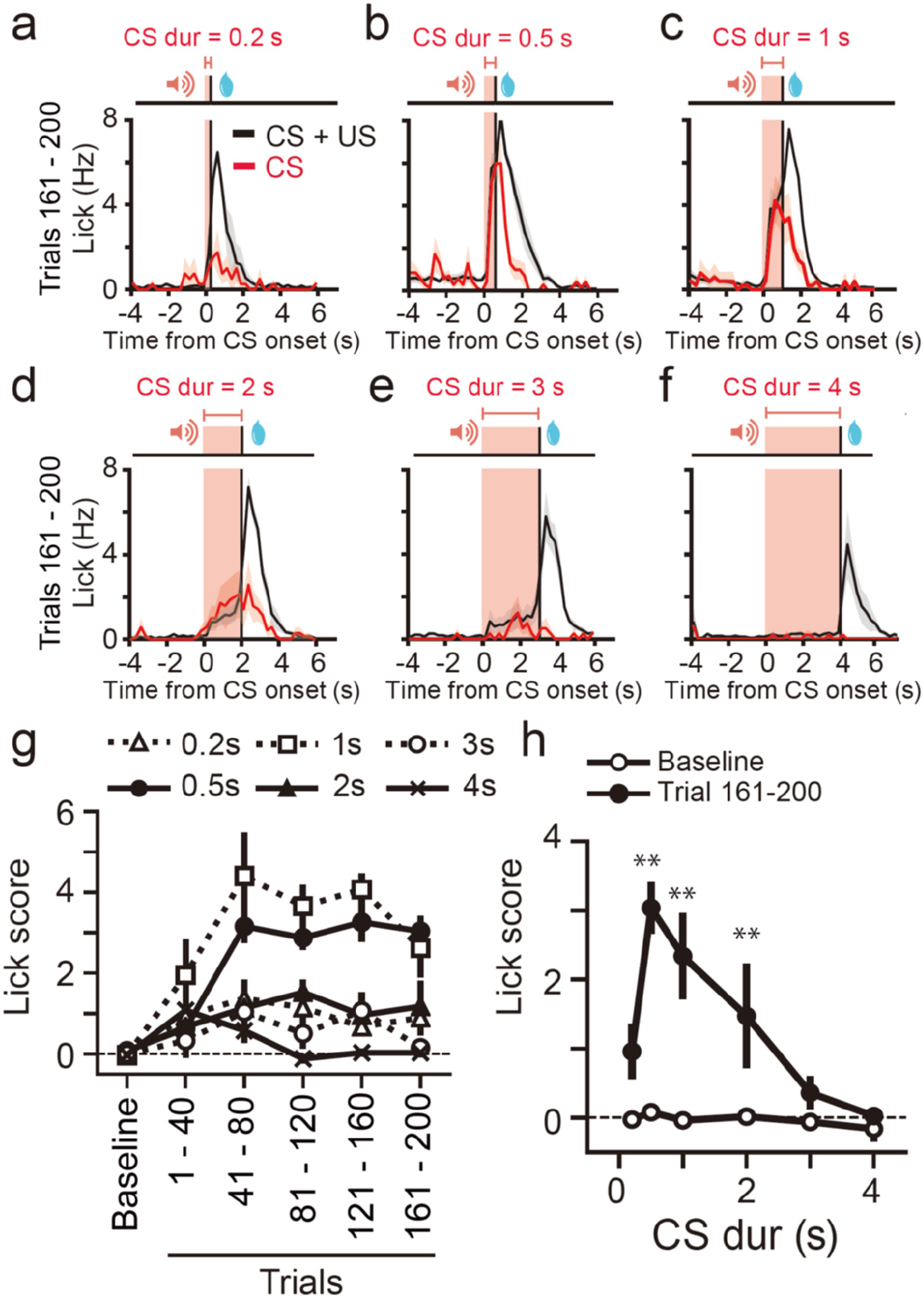
Conditioning with various CS durations. a-f) Averaged PSTHs of the licking responses with CS durations of 0.2 s (a, *n* = 4 mice), 0.5 s (b, *n* = 7 mice), 1 s (c, *n* = 5 mice), 2 s (d, *n* = 7 mice), 3 s (e, *n* = 4 mice), or 4 s (f, *n* = 5 mice) during trials 161–200. The plot in (b) is the same as that in the bottom of Fig. 1f and Fig. 2d. Red shades indicate the period of CS presentation. g) The lick scores for each condition plotted against time. Each symbol represents the CS duration. h) The effect of CS duration on conditioning. The averaged lick scores in the baseline session (white circle) and CS-only trials included during conditioning trials 161–200 (black circle) were plotted against CS durations. Two-way ANOVA with the test period as the repeated measurement factor. ** *P* < 0.01.

**Figure 3-Figure supplement-1.**
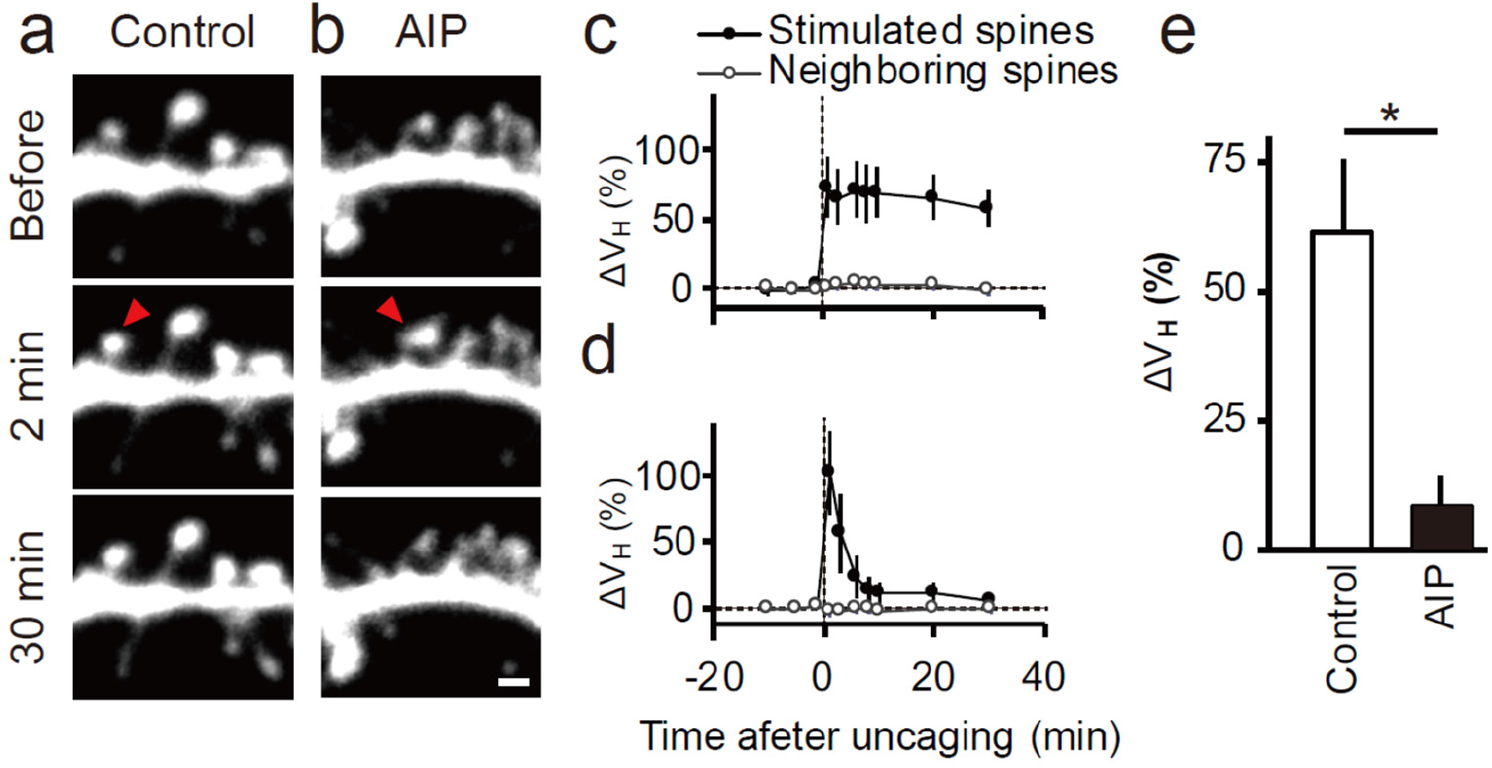
Blockade of spine enlargement by a CaMKII inhibitory peptide (AIP). a, b) Enlargement of the dendritic spines of the control vector-expressing SPNs (a) or AIP-expressing SPNs (b) in the NAc induced by two-photon uncaging of caged glutamate in a Mg-free ACSF. Red arrow heads indicate the targeted spines. The scale bar indicates 1 μm. c, d) Traces of the volume change of the dendritic spines with two-photon uncaging of caged-glutamate (stimulated spine) or spines located neighboring to the stimulated spines for control (c) and AIP (d). e) Averaged volume change 20 to 30 min after the stimulation. *n* = 6 dendrites, including 24 spines from 4 mice for AIP and *n* = 6 dendrites, including 23 spines from 3 mice for controls, * *P* < 0.05, t-test.

**Figure 3-Figure supplement-2.**
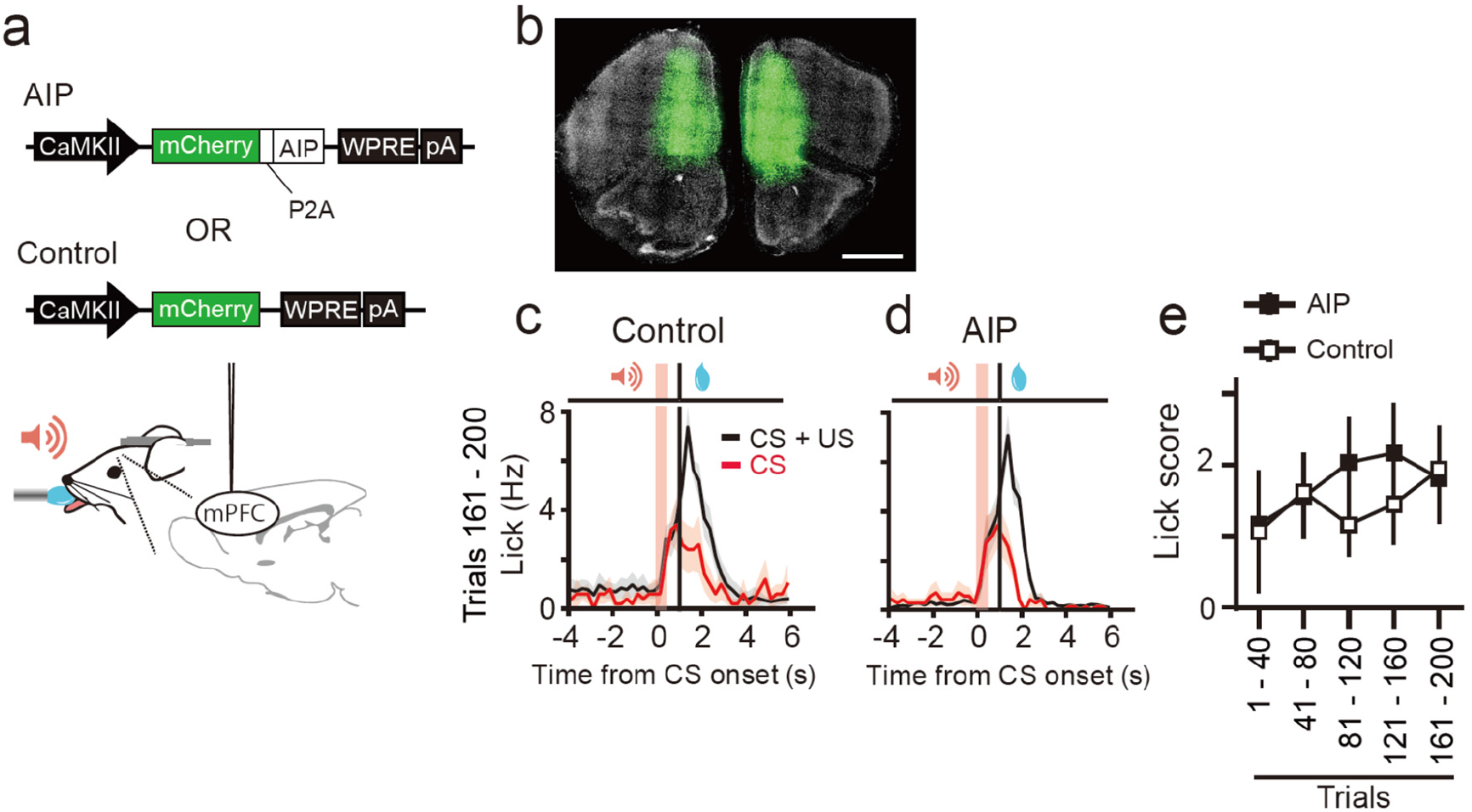
The effect of a CaMKII inhibitory peptide (AIP) on the mPFC on conditioning. a) Viral constructs and schematics of the injection. b) Confocal images of mCherry fluorescence (green) and DAPI (white) from a coronal slice including the PFC. The scale bar indicates 1 mm. c, d) Averaged PSTHs of the licking responses in CS + US (black) and CS (red) conditionings from mice injected with control (d, *n* = 5 mice) or AIP (c, *n* = 7 mice). e) The peak lick scores plotted against time for the conditions indicated.

**Figure 4-Figure supplement-1.**
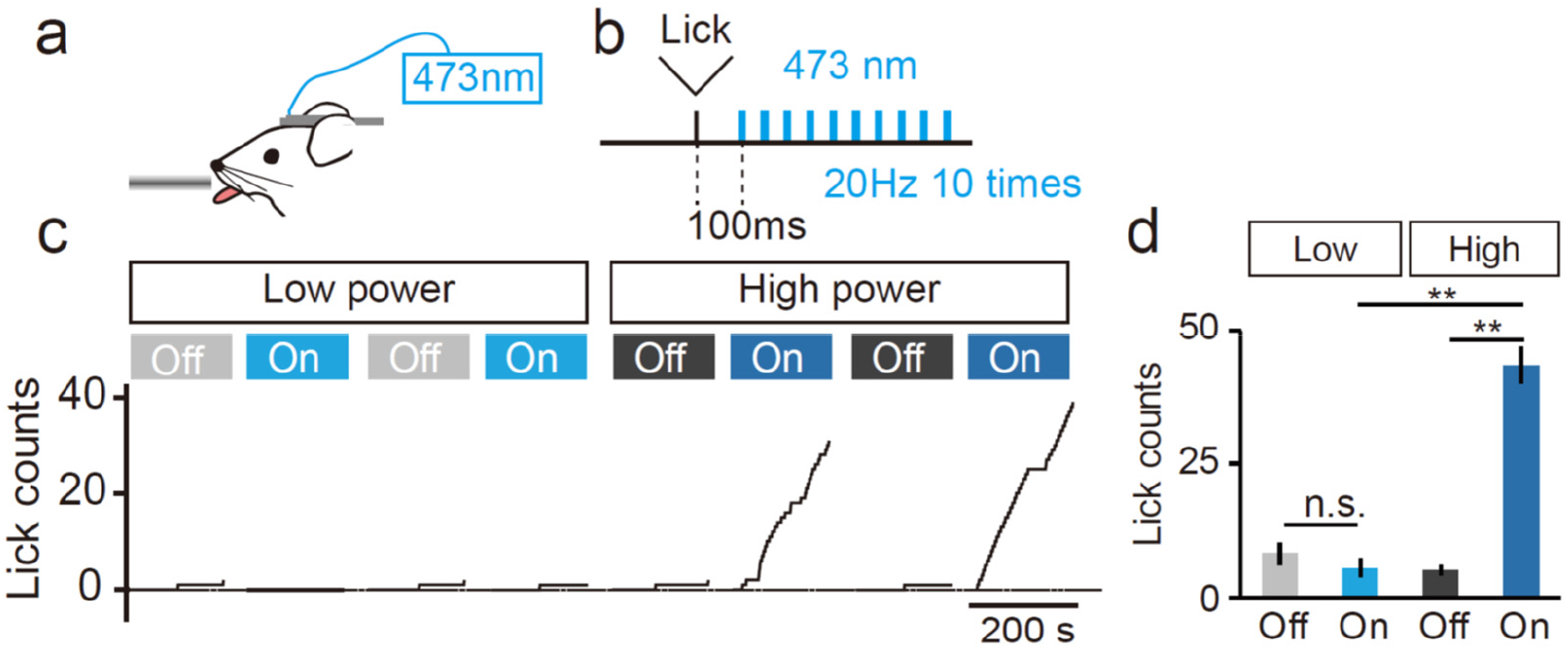
Operant conditioning with stimulation of the axon terminals from the BLA to the NAc. a, b) Operant conditioning using stimulation of the axon terminals from the BLA. The lick port detected licking to trigger blue laser stimulation (20 Hz, 10 times, 5 ms pulse width) with a delay of 100 ms (b). No water was given during this operant protocol. Laser power was adjusted to 1~3 mW for a low power condition and 5~15 mW for high power condition. c) Representative traces of cumulative licking counts during low power (2.5 mW) and high power (5 mW) conditioning. Conditions with laser off and on were alternated. d) Averaged total lick counts for the condition indicated. ** *P* < 0.01, n.s., not significant. One-way repeated measure ANOVA and Bonferroni post-hoc test.

**Figure 4- Figure supplement-2.**
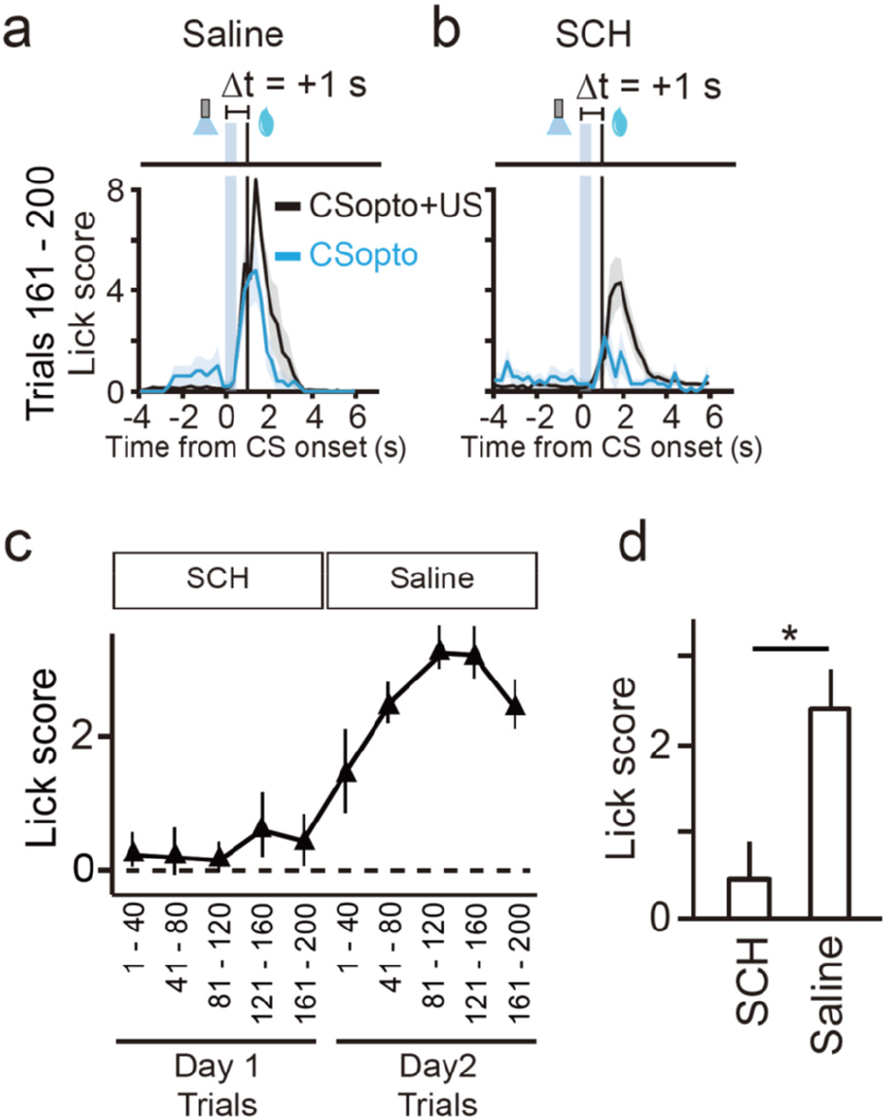
Effects of a dopamine D1 receptor blocker on Pavlovian conditioning with CSopto. a, b) Averaged PSTHs of the licking responses during the last 20% of the trials (161–200, bottom) for a dopamine D1 receptor blocker (SCH23390, 0.25 or 0.5 mg/kg, i.p) (a) or saline (b). On day 1, mice received intraperitoneal injections of SCH23390 before conditioning with CSopto with a delay of +1 s as in Fig. 4. The next day, the mice were administered saline and the conditioning was repeated. *n* = 5. c) Lick scores plotted against time. d) The effect of a dopamine D1 receptor blocker. Averaged lick scores from 4 CS-only trials during trials 161–200 were plotted for each condition. * *P* < 0.05, paired t-test, *n* = 5 mice.

**Figure 5- figure supplement-1.**
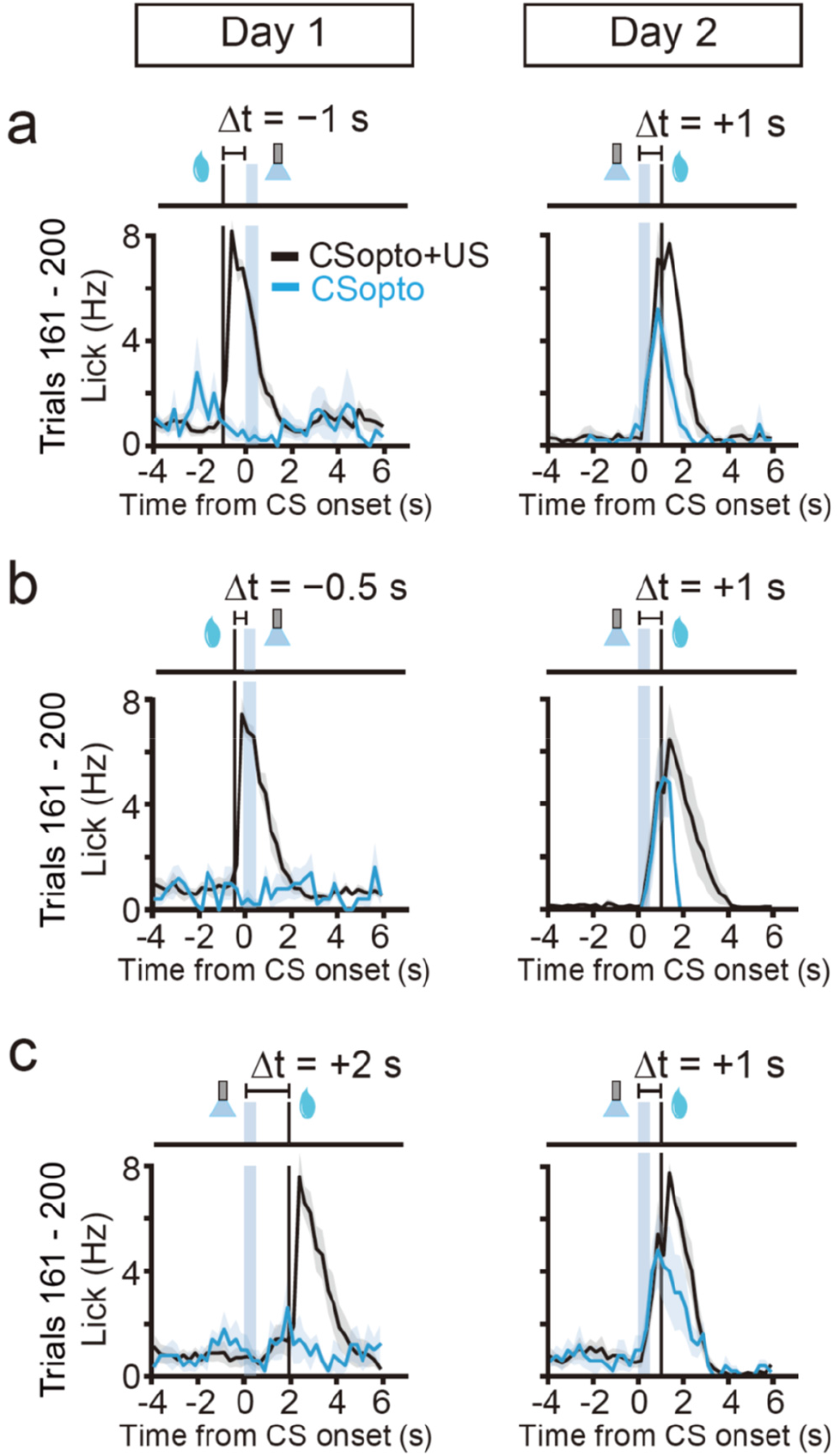
Positive controls for conditioning with CSopto. a, b, c) Averaged PSTHs of the licking responses during the last 20% of the trials (161–200, bottom). Mice allocated to Δt = −1 s (a, *n* = 5 mice, left), Δt = −0.5 s (b, *n* = 5 mice, left), and Δt = 2 s (c, *n* = 5 mice, left) on day 1 were conditioned with Δt = +1 s on day 2 (right).

## References

Abrams TW, Kandel ER (1988) Is Contiguity Detection in Classical-Conditioning a System or a Cellular Property – Learning in Aplysia Suggests a Possible Molecular Site. Trends in neurosciences 11: 128–135.

Bakker B (2002) Reinforcement learning with long short-term memory. In: Advances in neural information processing systems, pp 1475–1482.

Bangasser DA, Waxler DE, Santollo J, Shors TJ (2006) Trace conditioning and the hippocampus: the importance of contiguity. J Neurosci 26: 8702–8706.

Boice R, Denny MR (1965) The conditioned licking response in rats as a function of the CS-UCS interval. Psychonomic Science 3: 93–94.

Brea J, Gaal AT, Urbanczik R, Senn W (2016) Prospective Coding by Spiking Neurons. PLoS Comput Biol 12: e1005003.

Brzosko Z, Schultz W, Paulsen O (2015) Retroactive modulation of spike timing-dependent plasticity by dopamine. Elife 4: e09685.

Cassenaer S, Laurent G (2012) Conditional modulation of spike-timing-dependent plasticity for olfactory learning. Nature 482: 47–52.

Day JJ, Wheeler RA, Roitman MF, Carelli RM (2006) Nucleus accumbens neurons encode Pavlovian approach behaviors: evidence from an autoshaping paradigm. Eur J Neurosci 23: 1341–1351.

Ellis-Davies GC, Matsuzaki M, Paukert M, Kasai H, Bergles DE (2007) 4-Carboxymethoxy-5,7-dinitroindolinyl-Glu: an improved caged glutamate for expeditious ultraviolet and two-photon photolysis in brain slices. J Neurosci 27: 6601–6604.

Eshel N, Bukwich M, Rao V, Hemmelder V, Tian J, Uchida N (2015) Arithmetic and local circuitry underlying dopamine prediction errors. Nature 525: 243–246.

Fisher SD, Robertson PB, Black MJ, Redgrave P, Sagar MA, Abraham WC, Reynolds JNJ (2017) Reinforcement determines the timing dependence of corticostriatal synaptic plasticity in vivo. Nat Commun 8: 334.

Fremaux N, Gerstner W (2015) Neuromodulated Spike-Timing-Dependent Plasticity, and Theory of Three-Factor Learning Rules. Front Neural Circuits 9: 85.

Gallagher M, Graham PW, Holland PC (1990) The amygdala central nucleus and appetitive Pavlovian conditioning: lesions impair one class of conditioned behavior. J Neurosci 10: 1906–1911.

Gerstner W, Lehmann M, Liakoni V, Corneil D, Brea J (2018) Eligibility Traces and Plasticity on Behavioral Time Scales: Experimental Support of NeoHebbian Three-Factor Learning Rules. Front Neural Circuits 12: 53.

Gore F, Schwartz EC, Brangers BC, Aladi S, Stujenske JM, Likhtik E, Russo MJ, Gordon JA, Salzman CD, Axel R (2015) Neural Representations of Unconditioned Stimuli in Basolateral Amygdala Mediate Innate and Learned Responses. Cell 162: 134–145.

Grieger JC, Choi VW, Samulski RJ (2006) Production and characterization of adeno-associated viral vectors. Nat Protoc 1: 1412–1428.

Hawkins RD, Carew TJ, Kandel ER (1986) Effects of interstimulus interval and contingency on classical conditioning of the Aplysia siphon withdrawal reflex. J Neurosci 6: 1695–1701.

Hayar A, Bryant JL, Boughter JD, Heck DH (2006) A low-cost solution to measure mouse licking in an electrophysiological setup with a standard analog-to-digital converter. J Neurosci Methods 153: 203–207.

He K, Huertas M, Hong SZ, Tie X, Hell JW, Shouval H, Kirkwood A (2015) Distinct Eligibility Traces for LTP and LTD in Cortical Synapses. Neuron 88: 528–538.

Hebb DO (1949) The organization of behavior: A neuropsychological theory: Psychology Press.

Heys JG, Dombeck DA (2018) Evidence for a subcircuit in medial entorhinal cortex representing elapsed time during immobility. Nat Neurosci 21: 1574–1582.

Hikida T, Kimura K, Wada N, Funabiki K, Nakanishi S (2010) Distinct roles of synaptic transmission in direct and indirect striatal pathways to reward and aversive behavior. Neuron 66: 896–907.

Holland PC (1980) CS–US interval as a determinant of the form of Pavlovian appetitive conditioned responses. J Exp Psychol: Animal Behavior Processes 6: 155.

Ito I, Ong RC, Raman B, Stopfer M (2008) Sparse odor representation and olfactory learning. Nat Neurosci 11: 1177–1184.

Jocham G, Brodersen KH, Constantinescu AO, Kahn MC, Ianni AM, Walton ME, Rushworth MF, Behrens TE (2016) Reward-Guided Learning with and without Causal Attribution. Neuron 90: 177–190.

Kelley AE, Smith-Roe SL, Holahan MR (1997) Response-reinforcement learning is dependent on N-methyl-D-aspartate receptor activation in the nucleus accumbens core. Proc Natl Acad Sci U S A 94: 12174–12179.

Maren S, Fanselow MS (1996) The amygdala and fear conditioning: has the nut been cracked? Neuron 16: 237–240.

Mariath HA (1985) Operant-Conditioning in Drosophila-Melanogaster Wild-Type and Learning Mutants with Defects in the Cyclic-Amp Metabolism. Journal of insect physiology 31: 779–787.

Matsuzaki M, Honkura N, Ellis-Davies GC, Kasai H (2004) Structural basis of long-term potentiation in single dendritic spines. Nature 429: 761–766.

Menegas W, Babayan BM, Uchida N, Watabe-Uchida M (2017) Opposite initialization to novel cues in dopamine signaling in ventral and posterior striatum in mice. Elife 6: e21886.

Moe RO, Nordgreen J, Janczak AM, Spruijt BM, Zanella AJ, Bakken M (2009) Trace classical conditioning as an approach to the study of reward-related behaviour in laying hens: A methodological study. Applied Animal Behaviour Science 121: 171–178.

Murakoshi H, Shin ME, Parra-Bueno P, Szatmari EM, Shibata ACE, Yasuda R (2017) Kinetics of Endogenous CaMKII Required for Synaptic Plasticity Revealed by Optogenetic Kinase Inhibitor. Neuron 94: 37–47.

Otis JM, Namboodiri VM, Matan AM, Voets ES, Mohorn EP, Kosyk O, McHenry JA, Robinson JE, Resendez SL, Rossi MA, Stuber GD (2017) Prefrontal cortex output circuits guide reward seeking through divergent cue encoding. Nature 543: 103–107.

Pavlov IP (1927) Conditioned reflexes; an investigation of the physiological activity of the cerebral cortex. London: Oxford University Press: Humphrey Milford.

Reed SJ, Lafferty CK, Mendoza JA, Yang AK, Davidson TJ, Grosenick L, Deisseroth K, Britt JP (2018) Coordinated Reductions in Excitatory Input to the Nucleus Accumbens Underlie Food Consumption. Neuron 99: 1260–1273.

Rescorla RA (1988) Behavioral studies of Pavlovian conditioning. Annu Rev Neurosci 11: 329–352.

Rescorla RA, Holland PC (1982) Behavioral-Studies of Associative Learning in Animals. Annual Review of Psychology 33: 265–308.

Rombouts JO, Bohte SM, Roelfsema PR (2015) How Attention Can Create Synaptic Tags for the Learning of Working Memories in Sequential Tasks. Plos Computational Biology 11: e1004060.

Roseberry TK, Lee AM, Lalive AL, Wilbrecht L, Bonci A, Kreitzer AC (2016) Cell-Type-Specific Control of Brainstem Locomotor Circuits by Basal Ganglia. Cell 164: 526–537.

Rossi MA, Li HE, Lu D, Kim IH, Bartholomew RA, Gaidis E, Barter JW, Kim N, Cai MT, Soderling SH, Yin HH (2016) A GABAergic nigrotectal pathway for coordination of drinking behavior. Nat Neurosci 19: 742–748.

Schultz W, Dayan P, Montague PR (1997) A neural substrate of prediction and reward. Science 275: 1593–1599.

Sippy T, Lapray D, Crochet S, Petersen CC (2015) Cell-Type-Specific Sensorimotor Processing in Striatal Projection Neurons during Goal-Directed Behavior. Neuron 88: 298–305.

Smith-Roe SL, Kelley AE (2000) Coincident activation of NMDA and dopamine D1 receptors within the nucleus accumbens core is required for appetitive instrumental learning. J Neurosci 20: 7737–7742.

Stuber GD, Sparta DR, Stamatakis AM, van Leeuwen WA, Hardjoprajitno JE, Cho S, Tye KM, Kempadoo KA, Zhang F, Deisseroth K, Bonci A (2011) Excitatory transmission from the amygdala to nucleus accumbens facilitates reward seeking. Nature 475: 377–380.

Sutton RS, Barto AG (1998) Introduction to reinforcement learning: MIT press Cambridge.

Suvrathan A, Payne HL, Raymond JL (2016) Timing Rules for Synaptic Plasticity Matched to Behavioral Function. Neuron 92: 959–967.

Suzuki E, Nakayama M (2011) VCre/VloxP and SCre/SloxP: new site-specific recombination systems for genome engineering. Nucleic Acids Res 39: e49.

Thorndike EL (1911) Animal intelligence: Experimental studies: Macmillan.

Tolman EC (1948) Cognitive maps in rats and men. Psychol Rev 55: 189–208.

Tully T, Quinn WG (1985) Classical conditioning and retention in normal and mutantDrosophila melanogaster. Journal of Comparative Physiology A 157: 263–277.

Tye KM, Stuber GD, de Ridder B, Bonci A, Janak PH (2008) Rapid strengthening of thalamo-amygdala synapses mediates cue-reward learning. Nature 453: 1253–1257.

Wieland S, Schindler S, Huber C, Kohr G, Oswald MJ, Kelsch W (2015) Phasic Dopamine Modifies Sensory-Driven Output of Striatal Neurons through Synaptic Plasticity. J Neurosci 35: 9946–9956.

Wikenheiser AM, Schoenbaum G (2016) Over the river, through the woods: cognitive maps in the hippocampus and orbitofrontal cortex. Nat Rev Neurosci 17: 513–523.

Yagishita S, Hayashi-Takagi A, Ellis-Davies GC, Urakubo H, Ishii S, Kasai H (2014) A critical time window for dopamine actions on the structural plasticity of dendritic spines. Science 345: 1616–1620.

